# Chromatin-mediated alternative splicing regulates cocaine reward behavior

**DOI:** 10.1101/798009

**Authors:** Song-Jun Xu, Sonia I. Lombroso, Marco Carpenter, Dylan M. Marchione, Peter J. Hamilton, Carissa J. Lim, Rachel L Neve, Elizabeth A. Heller

**Affiliations:** Department of Systems Pharmacology and Translational Therapeutics, University of Pennsylvania, Philadelphia, PA, 19104, United States of America; Institute for Translational Medicine and Therapeutics, University of Pennsylvania, Philadelphia, PA, 19104, United States of America; Penn Epigenetics Institute, Perelman School of Medicine, University of Pennsylvania, Philadelphia, PA, 19104, United States of America; Department of Biochemistry and Biophysics, Perelman School of Medicine, University of Pennsylvania, Philadelphia, PA 19104, USA; Nash Family Department of Neuroscience, Friedman Brain Institute, Icahn School of Medicine at Mount Sinai, New York, NY, USA; Gene Delivery Technology Core, Massachusetts General Hospital, Boston, MA, USA

## Abstract

Alternative splicing is a key mechanism for neuronal gene regulation, and is grossly altered in mouse brain reward regions following investigator-administered cocaine. It is well established that cocaine epigenetically regulates transcription, yet mechanism(s) by which cocaine-induced epigenetic modifications regulate alternative splicing is largely unexplored. Our group and others have previously identified the histone modification, H3K36me3, as a putative splicing regulator. However, it has not yet been possible to establish the direct causal relevance of this modification to alternative splicing in brain or any other context. We found that mouse cocaine self-administration caused widespread alternative splicing, concomitant with enrichment of H3K36me3 at splice junctions. Differentially spliced genes were enriched in the motif for splice factor, Srsf11, which was both differentially spliced and enriched in H3K36me3. Epigenetic editing led us to conclude that H3K36me3 functions directly in alternative splicing of Srsf11, and that Set2 mediated H3K36me3 bidirectionally regulates cocaine intake.

## INTRODUCTION

Alternative splicing is a key mechanism for gene regulation in brain. Aberrant splicing is implicated in myriad neurological diseases, such as autism (Gonatopoulos-Pournatzis et al., 2018; Quesnel-Vallières et al., 2016; Voineagu et al., 2011), Rett syndrome (Cheng et al., 2017; Kriaucionis and Bird, 2004; Li et al., 2016), Huntington’s disease (Lin et al., 1993; Sathasivam et al., 2013; Wood, 2013), spinal muscular atrophy (Cartegni et al., 2006; Lorson et al., 1999; Parente and Corti, 2018; Xiong et al., 2015) and schizophrenia (Gandal et al., 2018; Glatt et al., 2011; Morikawa and Manabe, 2010; Nakata et al., 2009; Wu et al., 2012). However, this mechanism of gene regulation is understudied in neuropsychiatric diseases, including drug addiction. Persistence of cocaine-driven changes in isoform expression may underlie the chronic nature of addiction, analogous to permanent alternative isoform expression during neuronal differentiation (Schwartzentruber et al., 2018; Xu et al., 2017). It is well established that cocaine epigenetically regulates gene expression (Calipari et al., 2016; Cates et al., 2018; Damez-Werno et al., 2012, 2016; Feng et al., 2014; Hamilton et al., 2017; Heller et al., 2014, 2016; Maze et al., 2011; Taniguchi et al., 2012), yet mechanism(s) by which cocaine-induced epigenetic modifications regulate alternative splicing is largely unexplored.

Recent bioinformatic studies find that epigenetic features may be more important than gene sequence in differentiating splicing patterns (De Almeida et al., 2011; Bonev et al., 2017; Pajoro et al., 2017; Yuan et al., 2017; Zhou et al., 2012). Using agnostic classical statistical and machine-learning methods we previously found that histone H3 lysing 36 trimethylation (H3K36me3) is the most informative histone post-translational modification (hPTM) in predicting splicing events in the nucleus accumbens (NAc), a key brain reward region (Hu et al., 2017), and in multiple tissues across development (Hu et al., 2018). Furthermore, overexpression of the histone methyltransferase, SET domain containing 2 (Set2) (McDaniel and Strahl, 2017; Strahl et al., 2002), in human stem cells leads to splice-site enrichment of H3K36me3 and alternative exon exclusion at the fibroblast growth factor (*FGF2*) gene (Luco et al., 2010). Despite this progress in identifying H3K36me3 as a splicing regulator, it has not yet been possible to distinguish between the mere presence and causal relevance of this modification in alternative splicing in brain or any other context.

To elucidate the role of H3K36me3 in cocaine-induced alternative splicing, we first quantified concomitant splice-site enrichment of H3K36me3 and differential isoform expression in the mouse NAc following cocaine self-administration (SA). We found that alternative splicing serves as a key transcriptional mechanism in three brain reward regions in the context of volitional cocaine seeking, expanding on prior findings limited to investigator administered cocaine. Second, we showed that cocaine self-administration led to enrichment of H3K36me3 at splice junctions of differentially spliced genes in the NAc. Third, to validate that enrichment of H3K36me3 was causally linked to alternative splicing, we overexpressed Set2 in the NAc. Overexpression of Set2 and enrichment of H3K36me3 caused enhanced cocaine reward behavior, while pharmacological inhibition of Set2 attenuated the rewarding effects of cocaine. Global enrichment of H3K36me3 by either cocaine or Set2 led to alternative splicing of a common set of genes. We identified the splice factor, serine and arginine rich splicing factor 11 (Srsf11, also known as SRp54) (Gonatopoulos-Pournatzis et al., 2018), as highly enriched amongst both cocaine and Set2 spliced genes. Interestingly, Srsf11 was both differentially spliced and differentially enriched in H3K36me3 following either treatment. Finally, we applied targeted epigenetic editing to establish that H3K36me3 functions directly in alternative splicing of Srsf11, and found that this manipulation enhanced cocaine-reward behavior.

## RESULTS

### Alternative splicing and H3K36me3 enrichment in NAc following cocaine self-administration

Cocaine SA is a contingent paradigm for modeling addiction (Heilig et al., 2016), in which mice are trained to associate an operant response (e.g. spinning a wheel) with cocaine reward and reward delivery cues (light, tone). Mice also learn to discriminate between a cocaine-paired response and a control, saline-paired response. Control animals are placed in an identical experimental apparatus, but receive saline only. This complex paradigm recapitulates drug motivation and saliency, as well as drug/cue-associations, all of which are key features of the human disease (Sadri-Vakili et al., 2010; Schmidt and Pierce, 2010; Vassoler et al., 2013). To investigate alternative splicing and H3K36me3 enrichment in cocaine SA, mice (n=12) were trained to self-administer cocaine or saline (control) (Figure 1A). Measurement of drug infusions showed that mice infused cocaine during 21 2hr-SA sessions, and discriminated between the cocaine and saline-paired responses (Figure 1B, Supplement figure 1 A-C). For downstream biochemical analyses, mice were euthanized 24 hours after the last SA session, and tissue from brain reward regions was collected.

**Figure 1.**
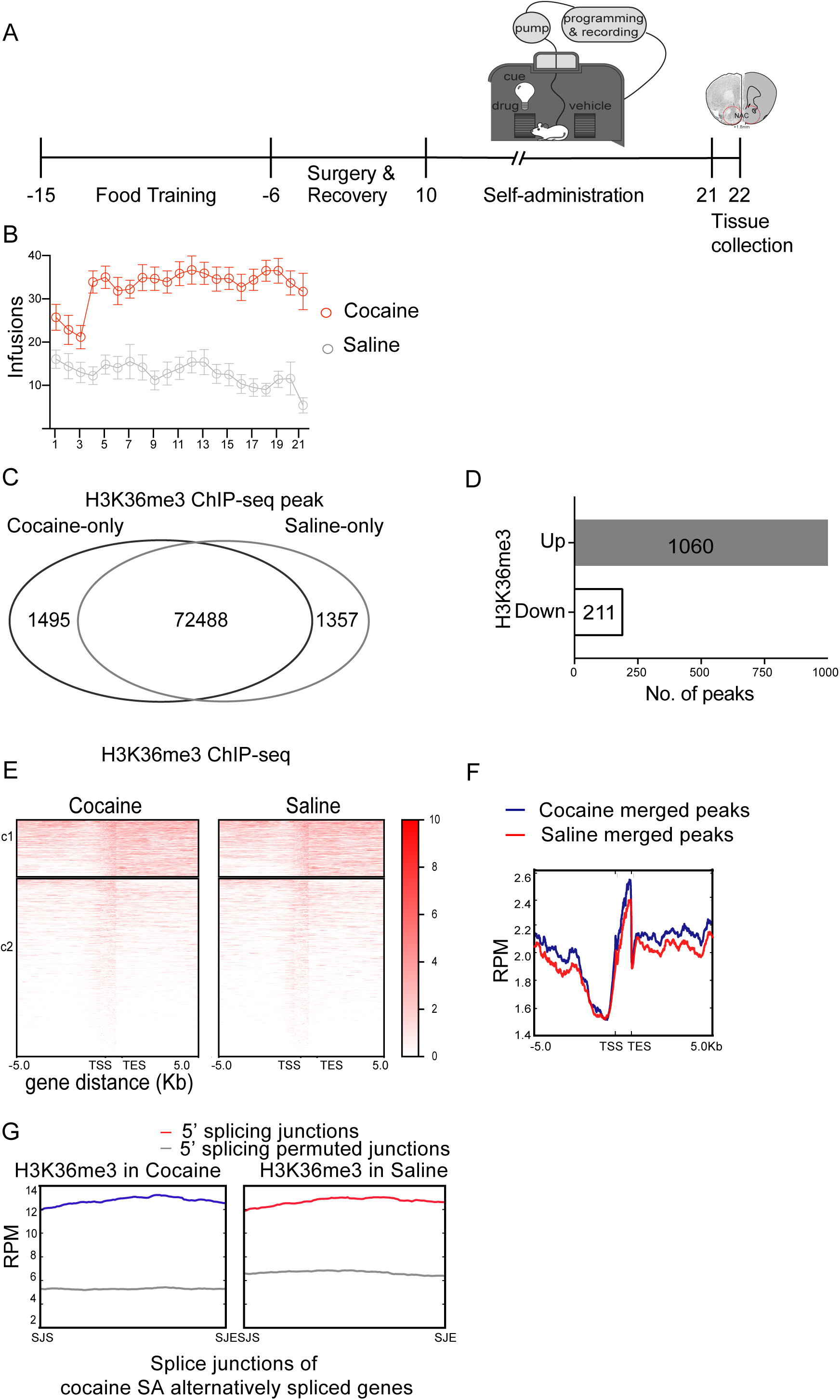
Alternative splicing and H3K36me3 enrichment in NAc following cocaine self-administration. (A) Schematic showing cocaine SA. Mice were food trained for 10 days prior to jugular catheterization surgery. Following recovery, mice (n=12/group) underwent 2hr, daily SA sessions for 21 days. Cocaine treated mice could self-administer either cocaine or saline, while saline treated controls could administer only saline. Following the last day of SA, mice were subjected to forced (home cage) abstinence for 1 day. NAc, VTA, and PFC tissue was then collected for downstream analysis. (B) Average cocaine or saline infusions (n=12 per group) per session for cocaine SA treatments. Mice showed a significant higher infusion rate for cocaine than saline in both treatments (two-way ANOVA repeated measure; Interaction effect F (20,440) =3.081, P<0.0001; drug effect F(1,22)=78.9, P<0.0001; session effect F (8.522,187.5) =2.298, P=0.0201). (C) Venn diagram showing cocaine-driven differential enrichment of H3K36me3 identified by ChIP-seq, compared to saline. Enrichment regions were counted based on Fold change > 1 or < −1 and FDR < 0.1. (D) Differential H3K36me3 enrichment after cocaine SA, compared to saline. Fold change > 1 or < −1 and FDR < 0.1. (E) H3K36me3 genic distribution after cocaine or saline SA. (F) Profile plot of H3K36me3 genome-wide (scaled gene body regions with 5kb up-/down-stream) distribution in cocaine SA and saline control. (G) Differential H3K36me3 enrichment at alternative splice junctions (+/-200bp) after cocaine (blue) or saline (red) treatment, compared to permuted junction sequences (grey). Splice junction start (SJS); splice junction end (SJE); reads per million (RPM).

To measure cocaine driven alternative splicing, we first applied rMATS splicing analysis (Shen et al., 2014) to RNA-sequencing data from the nucleus accumbens (NAc), prefrontal cortex (PFC) and ventral tegmental area (VTA), providing a comprehensive dataset of differential isoform expression in three brain reward regions (Supplement figure 1E-G). We next analyzed NAc RNA expression data with a pairwise comparison method with alternative splicing detection software MAJIQ (Norton et al., 2018). MAJIQ detects, quantifies, and visualizes local splicing variations (LSV), without dependence on existing isoform annotations. The relative abundance of each isoform is quantified as percent spliced in (PSI); ΔPSI quantifies the difference in relative isoform abundance between treatments. A pairwise comparison between treatments is more amenable to bioinformatic analysis of brain samples, which are heterogeneous both in tissue composition and behavioral endpoints, in that it extracts ΔPSI specific to between-group comparisons from that measured within-group. We identified 2730 genes that were differentially spliced in the NAc following cocaine SA (Supplement table 1).

In addition to alternative splicing, cocaine SA caused concomitant changes in global H3K36me3 enrichment, with approximately 2% of H3K36me3 peaks differentially enriched between saline and cocaine treated NAc (Figure 1C). More than 1000 H3K36me3 peaks (83.3%) were differentially enriched by cocaine SA, while 211 (16.6%) were de-enriched (Figure 1D). Cocaine SA did not lead to a gross redistribution of H3K36me3 (Figure 1C) nor a change in the genic distribution of H3K36me3 (Figures 1E-F). This is consistent with the fact that cocaine did not change expression of endogenous histone methyltransferase, Setd2 (Supplement figure 1D). To examine the correlation between alternative splicing events and H3K36me3 enrichment, we quantified H3K36me3 enrichment at splice junctions of alternatively spliced genes in NAc, following both saline and cocaine SA (Supplement table 1). As a control, we randomly permuted the genomic locations of the splicing junctions. We observed higher H3K36me3 enrichment at alternatively spliced junctions compared to permuted junction control, which exhibited a ‘flat-line’ like pattern (Figure 1G). This showed that H3K36me3 was specifically enriched at splice junctions of differentially spliced genes, suggesting that cocaine SA drives alternative splicing through regulation of H3K36me3 enrichment.

### Set2 overexpression in NAc regulated alternative splicing via H3K36me3 enrichment

We next sought to build on previous evidence that links Set2-dependent H3K36me3 to alternative splicing (De Almeida et al., 2011; Luco et al., 2010; Pajoro et al., 2017; Sessa et al., 2019; Xu et al., 2017; Yuan et al., 2017; Zhou et al., 2012). Although in higher eukaryotes multiple H3K36 methyltransferases catalyze H3K36me1 and H3K36me2, the yeast Set2 homolog is specific to H3K36me3 (McDaniel and Strahl, 2017; Sorenson et al., 2016; Strahl et al., 2002). We thus virally expressed Set2 to increase global H3K36me3 levels in NAc (Figure 2A, B). Expression of catalytically dead Set2(R195G) or GFP served as controls (Figure 2A). We validated global enrichment of H3K36me3 by western blot (Figure 2C) and quantitative histone mass spectrometry (Figure 2D). ChIP-seq analysis (n=3) showed genome-wide H3K36me3 enrichment after Set2 overexpression relative to Set2(R195G) (Figure 2E-H).

**Figure 2.**
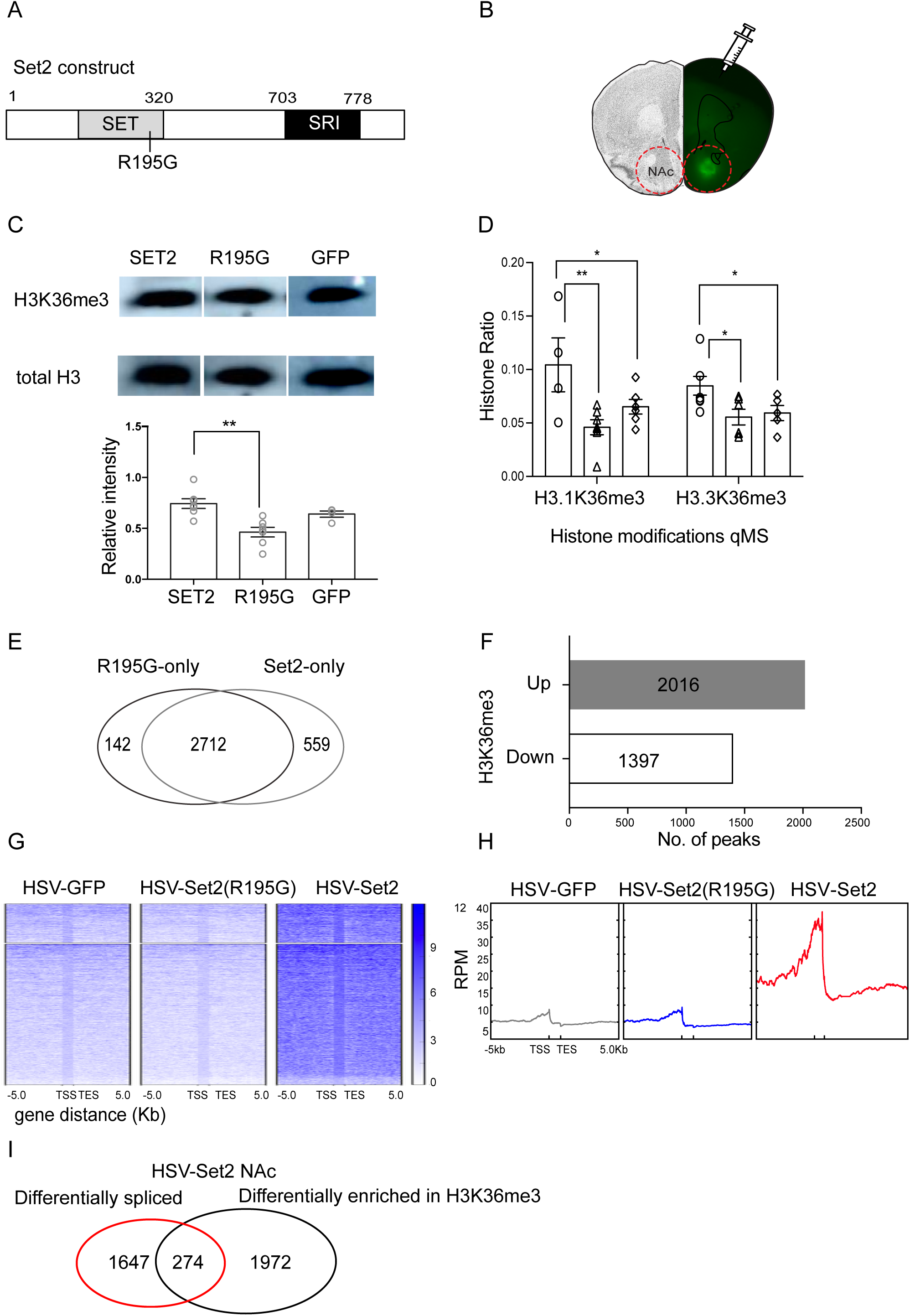
Set2 overexpression in NAc regulated alternative splicing via H3K36me3 enrichment. (A) Schematic of Set2 histone methyltransferase and catalytically dead Set2(R195G) control. SRI: Set2 Rpb1 interacting domain. (B) Representative image of HSV virus mediated GFP expression in NAc. (C) Representative western blot and quantification of H3K36me3 protein in HSV-Set2, HSV-Set2(R195G) or HSV-GFP injected NAc (One-way ANOVA with Tukey’s multiple comparison test. Main effect of virus, DF=19, **p<0.01). (D) Quantitative mass spectrometry of H3.1 and H3.3 K36me3 in HSV-Set2, HSV-Set2(R195G) and HSV-GFP injected NAc (two-way ANOVA; Interaction effect F(2,29) =1.0521, P=0.362; Histone location effect F(1,29)=0.411, P=0.526, virus effect F(1,456) =9.7348, P<0.001)). (E) Venn diagram Set2-mediated differential enrichment of H3K36me3 identified by ChIP-seq, compared to Set2(R195G). (F) Quantification of up and down regulated H3K36me3 peaks in HSV-Set2 compared to HSV-Set2(R195G) contro. Fold change > 1 or < −1 and FDR < 0.1. (G) Heatmap of H3K36me3 distribution on gene bodies with 5kb up- and down-stream regions across genome in HSV-GFP, HSV-Set2(R195G), HSV-Set2 viral injected NAc. (H) Profile plot of H3K36me3 distribution of HSV-GFP, HSV-Set2(R195G), and HSV-Set2 viral injected NAc. (I) Venn diagram showing overlap between genes both differentially enriched in H3K36me3 and differentiall spliced following Set2 overexpression, compared to Set2(R195G). Fisher exact test (FET) P<0.00001.

To elucidate how Set2-induced enrichment of H3K36me3 affects splicing, we analyzed alternative splicing events using MAJIQ, with a pairwise comparison approach (Norton et al., 2018). We identified 2841 differential splicing events between Set2 and Set2(R195G) treated NAc (ΔPSI > |0.1|, FDR < 0.05) (Supplement table 2). Set2 overexpression in NAc caused a subset of genes to be both differentially enriched in H3K36me3 and differentially spliced, compared to Set2(R195G) control (Figure 2I). This is consistent with our previous observation that H3K36me3 is enriched at alternative splice junctions following cocaine SA (Figure 1G).

Set2 overexpression also regulated gene expression in NAc (Supplement Figure 2A-C), consistent with the role of H3K36me3 in marking transcriptionally active loci (Strahl et al., 2002; Wilhelm et al., 2011). To understand the functional role of Set2 in cocaine SA, we compared differentially expressed genes (DEG) from Set2 and cocaine SA, and found a significant overlap between the two groups (Supplement figure 2D). We validated H3K36me3-mediated downregulation of transmembrane protein 25 (*Tmem25*) and upregulation of plexin A1 (*Plxna1*) (Supplement figure 2E) in both biological NAc replicates and Neuro2A (N2a) cells (Supplement figure 2F-G). Taken together, these results show Set2 overexpression increased global H3K36me3 enrichment in NAc affecting both gene transcription (Supplement figure 2A-C, 2E-G) and alternative splicing (Supplement table 2).

### Set2 bidirectionally regulates mouse cocaine reward behavior

Having established a potential role for Set2 in cocaine-induced alternative splicing, we next wanted to confirm its importance in cocaine reward. We first applied the cocaine conditioned place preference (CPP) paradigm (Mcclung, 2007) (Figure 3A), in which mice initially freely explore a two-sided chamber, to confirm a lack of innate preference for either side (pre-test). Mice are then trained to associate investigator-administered cocaine with exploration of only one side of the chamber, and saline with the other. After training, mice are again allowed to freely explore the two chambers (post-test). The difference in percentage of total time spent on the cocaine-paired side before (pre-test) and after (post-test) training quantifies CPP. Manipulations that enhance cocaine reward lead to increased time spent in the cocaine-paired side (Heller et al., 2016; Mcclung, 2007).

**Figure 3.**
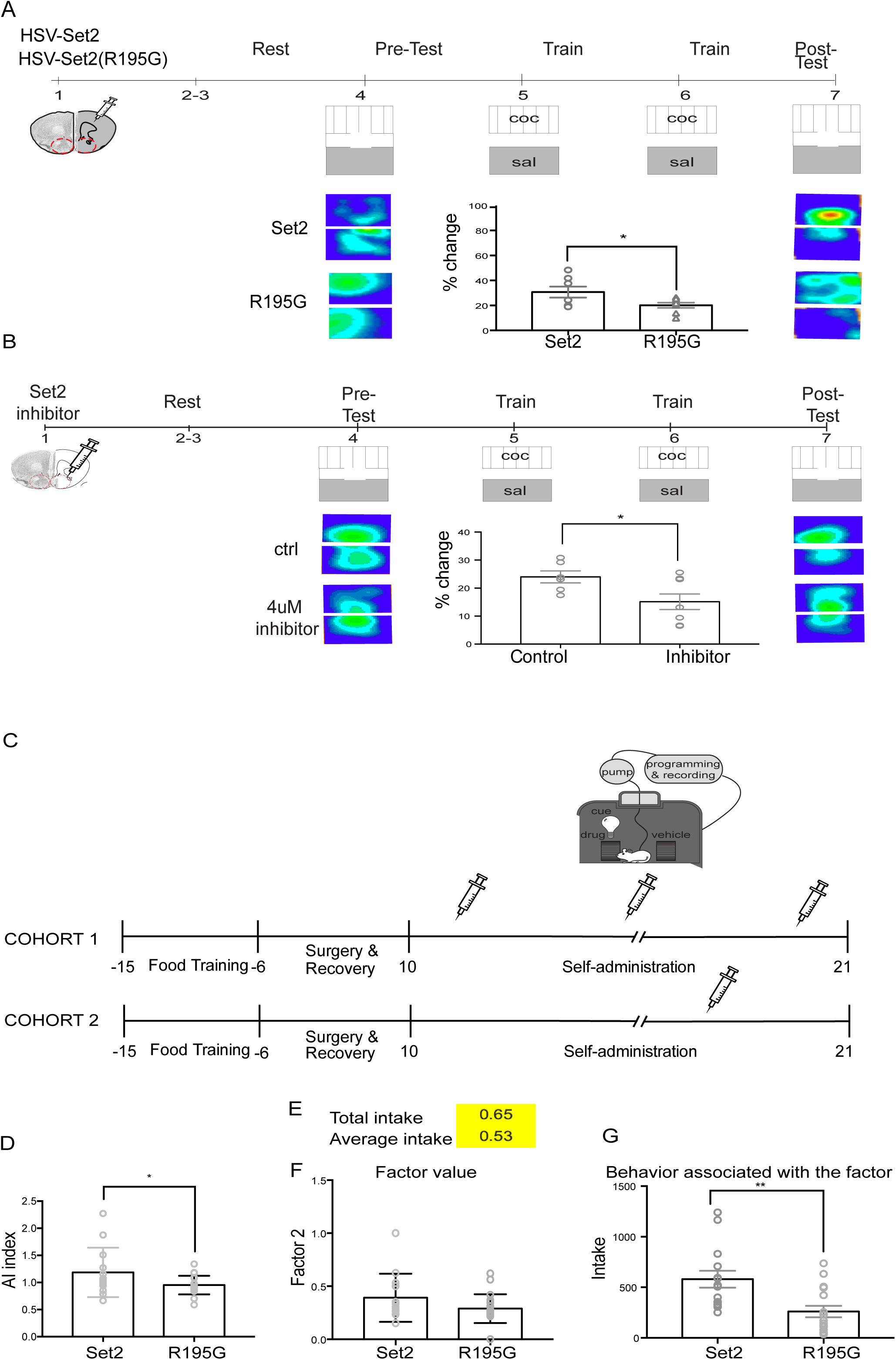
Set2 bidirectionally regulated mouse cocaine reward behavior. (A) Schematic for HSV-Set2 or HSV-Set2(R195G) viral injection followed by CPP behavioral test. Percentage change in time spent on cocaine-treated side by HSV-Set2 and -Set2(R195G) injected animals in CPP test (* p<0.05, student t-test). (B) Mice were injected intra-NAc with control (DMSO) or Set2 inhibitor (Bay598, 4 μM) followed by cocaine CPP. Set2 inhibition led to a decrease in percentage change in time spent on cocaine-treated side following training (* p<0.05, n=6-8, Student’s t-test). There was no difference (p=0.28) in total distance traveled between inhibitor- and control-treated mice, indicating that the reduction in CPP is not due to changes in locomotor behavior. (C) Schematic showing cocaine SA with Set2 viral injection. Mice were food trained for 10 days prior to jugular catheterization surgery. Following recovery, mice underwent 2hr, daily SA sessions for 21 days. HSV-Set2 virus were given at different time points between the two cohorts (indicated by syringes). Following the last day of SA, mice were subjected to forced (home cage) abstinence for 1 day. NAc tissue was then collected for downstream analysis. (D) Addiction index (AI) of factors that are most strongly associated with addiction-like behaviors. Set2 treated group showed significantly higher AI than Set2(R195G) group (Student T-test, * P<0.05). (E) Factor loading of factors 2 with SA behavior cocaine intake (yellow = positive). (F) Individual transformed data from cocaine total intake during SA following Set2 or Set2(R195G) NAc injection (Student T-test, non-significant). The transformed value was calculated to generate AI for each individual. (G) Individual cocaine total intake data presented for the behavior associated with factor 2 (Student T-test, ** P<0.01).

In our experiment, mice were injected intra-NAc with either HSV-Set2 or HSV-Set2(R195G), and subjected to cocaine CPP (Figure 3A). All mice receiving cocaine regardless of virus treatment showed an increase in percentage time spent in the cocaine-paired chamber (Figure 3A), indicating that virus treatment did not interfere with conditioning. Amongst cocaine treated mice, HSV-Set2 enhanced CPP compared to HSV-Set2(R195G), suggesting that H3K36me3 enrichment increases cocaine reward (Figure 3A, Supplement figure 3A-B). There was no difference in the total distance traveled between HSV-Set2 and - Set2(R195G) injected animals (Supplement figure 3A), indicating that Set2-enhanced cocaine reward is not due to changes in locomotor behavior. We further validated the relevance of H3K36me3 enrichment to cocaine CPP using an enzymatic inhibitor of Set2 (Figure 3B, Supplemental Figures 3B-D). We performed CPP after intracranial injection of the SET domain inhibitor, Bay598, to globally reduce H3K36me3, specifically in mouse NAc. Importantly, we found that Set2 inhibition reduced CPP compared to vehicle control (Figure 3B), suggesting that decreased H3K36me3 enrichment reduces the rewarding effects of cocaine. These data are especially significant given the need to identify novel targets that reverse the effects of drug exposure in order to treat addiction. Additionally, we found inhibition of Set2 enzymatic activity rescued its effects on cocaine CPP (Supplemental figures 3C) and had no effect on locomotor activity (Supplemental figure 3D). Lastly, to explore the role of H3K36me3 in bidirectional cocaine preference, we measured the correlation between global enrichment of H3K36me3 and cocaine CPP, finding a positive linear correlation (Supplement figure 3E).

Given our finding that global H3K36me3 enrichment enhanced cocaine CPP, we next tested the role of Set2 in cocaine SA, which evaluates animals’ motivation and cue discrimination, as well as drug intake (Sadri-Vakili et al., 2010; Schmidt and Pierce, 2010; Vassoler et al., 2013; Walker et al., 2018). Similar to human addicts, these features of drug behavior are variable across animal subjects. We used exploratory factor analysis to reduce multidimensional behavioral data to latent factors associated with interrelated behavioral traits (Figure 3E) (Walker et al., 2018). This approach discriminates between baseline individual differences in behavior and those driven by cocaine treatment (Supplement figure 3F). Using published criteria (Walker et al., 2018), we identified 3 factors that are associated with SA behavior and reflect important components of addiction: cocaine intake (Factor 2, Figures 3E-G), total paired responses (Factor 5, Supplement figures 3G-I), and discrimination between paired and unpaired responses (Factor 7, Supplement figures 3J-L). Based on these measures of addiction-related behaviors, we calculated a composite score, or Addiction Index (AI), for each animal (Walker et al., 2018). We found that animals with overexpression of Set2 in NAc had a higher AI (Figure 3D), compared to Set2(R195G) controls. Taken together, these behavioral data demonstrated that Set2 and global enrichment of H3K36me3 in NAc bidirectionally regulate cocaine reward behavior.

### Set2 and cocaine increased alternative splicing and H3K36me3 enrichment of Srsf11

We measured 1350 genes that were differentially spliced following both cocaine SA and Set2 overexpression, relative to saline and Set2(R195G), respectively (Figure 4A). Given this significant overlap, we hypothesized that the common differentially spliced genes were (1) regulated by the same splice factor(s) (2) causal to the Set2-mediated enhancement in cocaine reward behavior.

**Figure 4.**
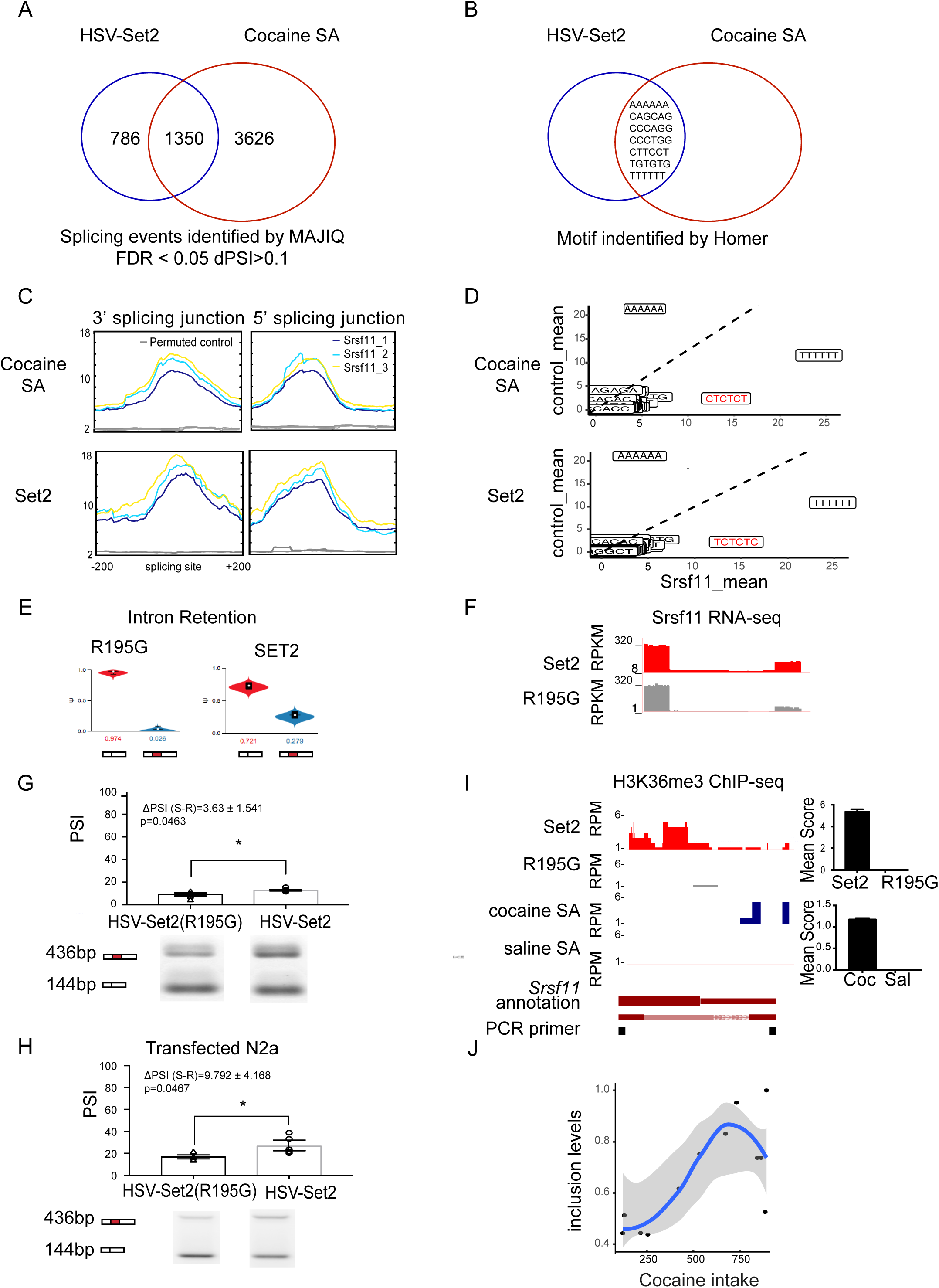
Set2 and cocaine regulated alternative splicing and H3K36me3 enrichment of *Srsf11*. (A) Venn diagram showing comparison of alternatively spliced genes in HSV-Set2 and cocaine SA treated groups (pairwise comparison using MAJIQ, FDR < 0.05 ΔPSI > 0.1; FET P < 0.00001). (B) Venn diagram showing shared 6-mer motifs that were independently identified in alternative splicing junctions of Set2 overexpression and cocaine SA treatments. (C) Profile plot of Srsf11 iCLIP-seq signal at alternative splicing junctions (top “bell-curve” like) and scrambled sequence control (bottom “flat-line” like) of cocaine SA and Set2 overexpression treatments. (D) Averaged z-score of Srsf11 binding frequency compared to control on splicing motifs identified in cocaine SA and Set2 splice junctions. (E) Violin plots of PSI values of Srsf11 intron retention alternative splicing event identified by MAJIQ in Set2(R195G) and Set2 overexpression in NAc. (F) RNA-seq track coverage at alternative splicing region in Set2(R195G) and Set2 overexpression in NAc. (G) Quantification for validating intron retention event in Set2(R195G) and Set2 overexpression in NAc with representative PCR blot (* p<0.05, student T-test). (H) Quantification for validating intron retention event in Set2 and -Set2(R195G) transfected N2a cells with representative PCR blot (* p<0.05, Student’s t-test). (I) H3K36me3 ChIP-seq coverage and quantification at Srsf11 alternative splice junction in Set2(R195G) and Set2 overexpression in NAc. (J) Positive linear correlation between NAc expression of Srsf11 alternative isoform and cocaine SA behavior (P<0.01, R^2^=0.73). Cocaine intake and Srsf11 alternative isoform expression was measured within-animal by cocaine self-administration followed by RNA-seq.

Application of the bioinformatic tools, Homer and MEME, identified 7 splice factor binding motifs common to both treatments (Figure 4B-C, Supplement figure 4A). The corresponding splice factors were identified using MEME-TOMTOM (Gupta et al., 2007) and published iCLIP-sequencing data sets from mouse neurons (Chen et al., 2018; Rodor et al., 2017; Takeuchi et al., 2018; Vuong et al., 2016) (Figure 4B, Supplement figure 4A,C). We focused on the motif for serine/arginine-rich splicing factor 11 (Srsf11), as it showed enrichment at the junction regions, with robust increases at exon start and end sites, and a ‘flat-line’-like pattern of enrichment at the permuted junction controls (Figure 4C). This computational data was further validated by the fact that the Srsf11 motifs identified in our dataset are also those most enriched by Srsf11 iCLIP (UCUCUC and CUCUCU) (Gonatopoulos-Pournatzis et al., 2018) (Figure 4D). 7 additional splice motifs were enriched in our dataset, including neuro-oncological ventral antigen 2 (*Nova2*), polypyrimidine tract binding protein 2 (*PTBP2*) (Saito et al., 2019), but the enrichment patterns at splice junctions were highly similar to the permuted controls, indicating that those splice factors were not enriched at splice junctions of alternatively spliced genes in Set2 or cocaine SA datasets (Supplement figure 4C).

The role of Srsf11 in driving differential splicing could not be linked to changes in its expression, as there were no differences in Srsf11 expression following cocaine SA (Supplement figures 4B, D) or Set2 overexpression (Figure 4F, Supplement figures 4B, F), relative to saline or Set2(R195G), respectively. Alternatively, Set2 overexpression and cocaine SA both led to alternative splicing of Srsf11 (Figure 4E, Supplement figure 4E), specifically intron retention (Supplement table 3). We biochemically validated these findings in a replicate cohort of HSV-Set2 or HSV-Set2(R195G) injected mice (Figure 4G) and Set2-transfected N2a cells (Figure 4H), using qPCR and PCR methods to measure gene expression (Supplement figure 4F) and alternative splicing (Figure 4G-H), respectively. PCR validation of alternative splicing was accomplished using a single pair of primers to amplify multiple isoforms; PSI was quantified as the relative abundance of each isoform in a single lane (Gonatopoulos-Pournatzis et al., 2018). Furthermore, both Set2 overexpression and cocaine SA caused differential enrichment of H3K36me3 at the Srsf11 splice junction (Figure 4I). Given our discovery that enrichment of H3K36me3 increased both cocaine reward behavior and intron inclusion of Srsf11, we hypothesized that this Srsf11 isoform was associated with cocaine reward behavior. Indeed, we measured a positive linear correlation between cocaine intake and Srsf11 inclusion levels (Figure 4J). Together, these findings suggested that both cocaine SA and Set2 drive a common differential splicing profile via alternative splicing of the splice factor, Srsf11. H3K36me3 enrichment at *Srsf11* splice junctions suggested a role for this hPTM in driving alternative splicing of this gene.

### Epigenetic editing of H3K36me3 was sufficient to drive alternative splicing of Srsf11 and cocaine reward behavior

Set2 overexpression in NAc established a causal relationship between H3K36me3 and splicing of Srsf11, but this experiment did not prove the direct causal relevance and sufficiency of H3K36me3 enrichment to Srsf11 splicing. To address this, we used nuclease-deficient Cas9 (dCas9) fused to Set2 for targeted epigenetic editing of H3K36me3. Cautiously, we noted previous data suggesting that when the Cas9-sgRNA complex is targeted intragenically to the non-template DNA strand it causes expression silencing, while complexes binding to the template DNA strand do not (Qi et al., 2013). We thus designed the *Srsf11* sgRNAs to target either the template (T1) or non-template (N2, N3) strands, with minimal predicted off-target mismatch (see methods) (Figure 5A). A non-targeting (NT) control sgRNA did not align to any genomic sequence.

**Figure 5.**
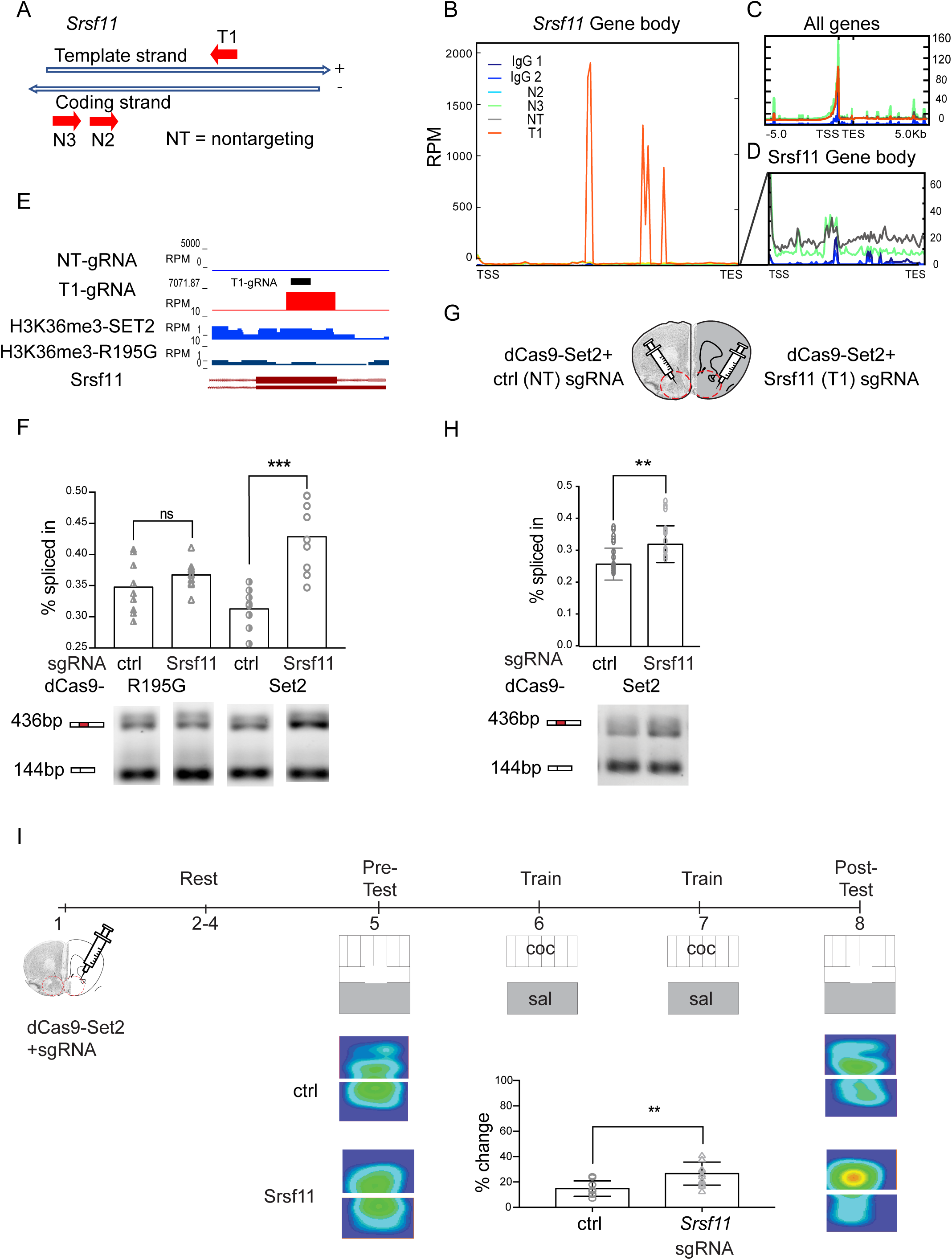
Epigenetic editing of H3K36me3 was sufficient to drive alternative splicing of Srsf11 and increase cocaine reward behavior. (A) Schematic of sgRNA designed to target H3K36me3 enriched region of *Srsf11*. (B) H3K36me3 enrichment at Srsf11 in N2a cells transfected with dCas9-Set2 and sgRNAs. Note that H3K36me3 enrichment following transfection with sgRNAs N2, N3 and NT at not above background, and are shown in panel D. T: Template, N: Non-template, NT: Non-targeting. (C) H3K36me3 enrichment at Srsf11 in N2a cells transfected with dCas9-Set2 and sgRNAs. (D) H3K36me3 enrichment at targeted Srsf11 gene body region of dCas9-Set2 with sgRNA N2, N3, NT and IgG controls. Note difference in RPM scale relative to panel B. (E) H3K36me3 CUT&RUN-seq track coverage showing H3K36me3 enrichment at sgRNA T1 targeted locus. Black bar: 20bp protospacer motif. (F) Quantification of Srsf11 alternative isoform expression in N2a cells transfected with dCas9-Set2 or dCas9-Set2(R195G) and sgRNA T1 or NT (ns, non-significant, *** p<0.001. One-way ANOVA with Tukey’s multiple comparison, P(dCas9-Set2-NT (ctrl) vs. dCas9-Set2-T) = 0.0007; P(dCas9-R195G-T vs. dCas9-R195G-NT(ctrl)) = 0.018). (G) Schematic showing in-vivo transfection of dCas9-SET2 with sgRNA T1 and NT in two hemispheres of NAc. (H) Quantification of Srsf11 alternative isoform expression in NAc expressing dCas9-Set2 and sgRNA T1 or NT (** p<0.01. Student T-test). (I) Timeline and schematic of NAc expression of dCas9-Set2 plus sgRNA T1 or NT, followed by CPP. Percentage change in time spent in cocaine-paired chamber following dCas9-Set2 plus sgRNA T1 or NT treatment (* p<0.05, **p<0.01 Student T-test P=0.0032).

We measured H3K36me3 enrichment using ‘Cleavage Under Targets and Release Using Nuclease’ (CUT&RUN)-sequencing (Skene and Henikoff, 2017; Skene et al., 2018) on N2a cells transfected with dCas9-Set2 and sgRNAs T1, N2, or N3 (Figure 5F). dCas9-Set2 plus sgRNA T1 led to a 200-fold increase of H3K36me3 enrichment compared to sgRNA non-targeting (NT) (Figure 5B), while sgRNAs N2 and N3 showed similar enrichment to NT (Figure 5D; note change in RPM scale). sgRNAs T1, N2, N3 and NT all showed similar global H3K36me3 distributions, suggesting negligible off-target effects (Figure 5C). Track coverage confirmed that the sgRNA T1-specific, H3K36me3 enriched region overlapped precisely with the sgRNA T1 photospacer motif, which was not observed for the control sgRNAs NT, N2 and N3 (Figure 5E). Using PCR, we found that dCas9-Set2 transfected with *Srsf11*-sgRNA-T1 increased Srsf11 intron retention (inclusion ratio) compared to control-sgRNA-NT control. No differences were observed between dCas9-Set2 plus control-sgRNA-NT and dCas9-Set2(R195G) plus *Srsf11*-sgRNAs-T1 or control-sgRNA-NT (Figure 5F). Together, these data showed that in N2a cells, targeted epigenetic editing of H3K36me3 at the splice junction is sufficient to drive alternative isoform expression of Srsf11 via increased intron retention.

To validate this result in vivo, we expressed dCas9-Set2 and *Srsf11*-sgRNA (T1) or control-sgRNA-NT in mouse NAc. Neuronal-specific expression was conferred by the human synapsin 1 gene promoter. For scientific rigor, we used within-animal controls wherein one hemisphere was injected with control-sgRNA-NT and the other hemisphere was injected with *Srsf11*-sgRNA (T1) (Figure 5G). We found that expression of dCas9-Set2 transfected with *Srsf11*-sgRNA-T1 with increased the Srsf11 alternative isoforms inclusion ratio compared to control-sgRNA-NT (Figure 5H).

Given that global Set2 overexpression enhanced cocaine reward behavior (Figure 3), we next tested the hypothesis that alternative splicing of Srsf11 is sufficient to increase cocaine conditioned place preference. We transfected dCas9-Set2 plus *Srsf11*-sgRNA-T1 or control-sgRNA-NT in mouse NAc and subjected animals to cocaine CPP (Figure 5I). dCas9-Set2 plus *Srsf11*-sgRNA-T1 caused an increase in cocaine conditioned place preference, as measured by percent change in time spent on the cocaine-paired side, compared to control-sgRNA-NT (Figure 5I). There was no difference in distance travelled between treatment groups (Supplement figure 5D). Taken together, these data show that targeted epigenetic editing H3K36me3 at the *Srsf11* splice junction was sufficient to drive alternative splicing and enhance cocaine reward behavior in mice.

## DISCUSSION

Alternative splicing is a key mechanism for gene regulation in brain. Neuron-specific isoform expression is essential to proper cell-type specification as shown from recent studies in neuronal development (Furlanis and Scheiffele, 2018; Saito et al., 2019; Schwartzentruber et al., 2018), disease (Gandal et al., 2018; de la Torre-Ubieta et al., 2016; Parikshak et al., 2016; Voineagu et al., 2011) and activity (Cates et al., 2018; Parikshak et al., 2016; Quesnel-Vallières et al., 2016). While many neurological diseases, including autism (Quesnel-Vallières et al., 2016; Voineagu et al., 2011), Rett syndrome (Cheng et al., 2017; Kriaucionis and Bird, 2004; Li et al., 2016), Huntington’s disease (Lin et al., 1993; Sathasivam et al., 2013; Wood, 2013), spinal muscular atrophy (Cartegni et al., 2006; Lorson et al., 1999; Parente and Corti, 2018; Xiong et al., 2015) and schizophrenia (Gandal et al., 2018; Glatt et al., 2011; Morikawa and Manabe, 2010; Nakata et al., 2009; Wu et al., 2012) have been linked to disruptions in alternative splicing, this mechanism of gene regulation is understudied in the context of drug abuse and addiction. Only one study thus far has reported that investigator administered cocaine results in greater changes in isoform expression than gene expression in the NAc (Feng et al., 2014). In the current study, we found that alternative splicing serves as a key transcriptional mechanism in three brain reward regions in the context of volitional cocaine seeking, expanding on prior findings limited to investigator administered cocaine. In addition, we showed that cocaine self-administration led to enrichment of H3K36me3 at splice junctions of differentially spliced genes in the NAc.

Persistence of cocaine-driven changes in isoform expression may underlie the chronic nature of addiction, analogous to permanent alternative isoform expression during cell-fate determination (Schwartzentruber et al., 2018; Xu et al., 2017). It is well established that cocaine epigenetically regulates gene expression (Calipari et al., 2016; Cates et al., 2018; Damez-Werno et al., 2012, 2016; Feng et al., 2014; Hamilton et al., 2017; Heller et al., 2014, 2016; Maze et al., 2011; Taniguchi et al., 2012), yet mechanism(s) by which cocaine-induced epigenetic modifications regulate alternative splicing is largely unexplored. Prior data indicate that a subset alternatively spliced genes after investigator-administered cocaine are enriched in the splice factor motif for A2BP1(Rbfox1/Fox-1), as well as the hPTM, H3K4me3 (Feng et al., 2014). In fact, co-immunoprecipitation of A2BP1 and H3K4me3 in NAc increases after cocaine administration (Feng et al., 2014).

To add to this nascent body of literature on chromatin-regulated alternative splicing in brain, we previously applied agnostic classical statistical and machine-learning methods to find that H3K36me3 is the most informative hPTM in predicting splicing events in the NAc (Hu et al., 2017) and multiple other neural tissues at several developmental timepoints (Hu et al., 2018). Analysis of NAc enrichment of 8 hPTMs in the exon flanking regions reveals that only H3K36me3 and H3K4me1 are differentially enriched with respect to skipped exon category and best predicted skipped exon events (Hu et al., 2017). Genome-wide mapping of histone modifications in many organisms has revealed their non-random distribution around exons, with H3K36me3 enriched in exonic regions compared to intronic regions (Ernst and Kellis, 2017; Hu et al., 2017; Leung et al., 2019; Luco et al., 2010; Meers et al., 2017; Xu et al., 2017). A recent study in human stem cells finds that H3K36me3 regulates alternative splicing events and are involved in nonsense-mediated mRNA decay of BRCA1-associated RING domain protein 1 (BARD1), which is crucial to activate Ataxia telangiectasia mutated (ATM) and ATM and RAD3-related (ATR) pathway and maintain cell pluripotency state (Xu et al., 2017). This study reveals that epigenetic features play a dominant role in decoding alternative splicing patterns, suggesting a key function of hPTMs in comprehensively understanding splicing machinery (Xu et al., 2017). In addition, specific hPTMS are strongly associated with different splicing types. H3K36me3 marks splicing related exons as shown in studies profiling H3K36me3 enrichment as lower in alternative splicing exons than constitutive exons (Hu et al., 2017; Kolasinska-Zwierz et al., 2009; Luco et al., 2010; Sessa et al., 2019; Wilhelm et al., 2011). H3K36me3 also facilitates efficient mRNA splicing through recruitment of an “adapter protein” to stabilize splice factors and regulate proper co-transcriptional splicesome assembly (Leung et al., 2019). This alternative splicing related enrichment is highly evolutionarily conserved between humans and mice (Kolasinska-Zwierz et al., 2009; Wilhelm et al., 2011).

We expanded upon these bioinformatic analyses with an experimental manipulation of Set2, finding that enrichment of H3K36me3 in NAc by Set2 overexpression caused changes in global alternative splicing. We hypothesized that H3K36me3-mediated alternative splicing is an underlying mechanism for cocaine reward behavior. We found a significant overlap in alternatively spliced genes following Set2 and cocaine SA treatments. De novo motif search revealed 7 splice motifs common to both treatments, suggesting that the increased cocaine preference following Set2 treatment is regulated, in part, by the same splice machinery as cocaine SA. By combining de novo motif search and CLIP-seq data from neurons, we identified Srsf11 as a putative splice factor regulating splicing across treatments. Given the recently discovered role of Srsf11 in microexon expression in neurons (Gonatopoulos-Pournatzis et al., 2018), it will be interesting to analyze microexon expression and splicing in our datasets, especially following dCas9-Set2 driven splicing of Srsf11. Furthermore, we expect that additional splice factors are involved in cocaine driven alternative splicing, such as A2BP1, the motif for which was identified in our datasets and is enriched in spliced genes following investigator administered cocaine (Feng et al., 2014). In identifying key splicing factors that regulate cocaine driven alternative splicing, we were limited to available database of splicing factor motifs and CLIP-seq data that matched our species and tissue type of interest. As more comprehensive CLIP-seq data become available, it will be possible to identify additional components of the splice machinery relevant in cocaine and chromatin-mediated alternative splicing.

Having identified Srsf11 as a likely candidate for the regulation of Set2 and cocaine-driven alternative splicing, we sought to analyze its mode of regulation in these contexts. Interestingly, while *Srsf11* gene expression was unchanged, both cocaine and Set2 caused increased intron retention in this transcript and enrichment of H3K36me3 at the relevant splice junction. Notably, the differential Srsf11 isoforms are associated with functional differences. The alternative inclusion isoform (ENSMUST00000152274) is translated into a 423aa coding protein (ENSMUSP00000127239) while the exclusion isoform (ENSMUST00000126716) is degraded by nonsense medicated mRNA decay (ENSMUSP00000114370). Increasing intron retention is a possible post-translational mechanism for overall increased expression of the Srsf11 protein.

We found that epigenetic editing using dCas9-Set2 was sufficient to recapitulate endogenous enrichment of H3K36me3 at the splice junction of *Srsf11* and drive alternative splicing both in vitro and in vivo. Our results indicate that H3K36me3 can directly regulate alternative splicing without affecting expression. However, given that H3K36me3 is associated with active transcription, the presence of this mark on a splice factor gene could indirectly regulate splicing. This mechanism can be further probed by identification of the splice factor genes differentially expressed following Set2 overexpression. Chromatin modifications may directly regulate splicing in specific cellular contexts, and may do so via recruitment of splice factors/adapter, or regulation of the kinetics of RNAPolII. Evidence for direct recruitment of an adapter system by H3K36me3 is demonstrated at the fibroblast growth factor (*FGFR2*) gene during cell differentiation (Luco et al., 2010). This adaptor system consisting of hPTM (H3K36me3), a chromatin-binding protein (mortality factor 4 like 1, MRG15) and a splicing regulator (polypyrimidine tract binding protein 1, PTB), regulates FGFR2 alternative splicing (Luco et al., 2010). However, in the prior study Set2 overexpression increased global H3K36me3, which makes it difficult to distinguish direct function of this hPTM in splicing FGFR2 from pleiotropic effects. To examine the direct causal relevance of H3K36me3 in splicing Srsf11 in NAc we took an epigenetic editing approach. dCas9-Set2 directed to Srsf11 recapitulated endogenous H3K36me3 enrichment at a single splice junction and drove splicing of this gene. Further studies will be aimed at distinguishing between the direct recruitment and kinetic models of splicing regulation by H3K36me3.

Epigenetic editing in brain has seen major advances in its application to gene activation and silencing (Lee et al., 2018; Liu et al., 2018; Zheng et al., 2018). For example, targeted dCas9-Tet1 to CGG repeats in gene fragile X mental retardation 1 (*FMR1)* is sufficient to reverse the hypermethylation at this locus and sustain expression of fragile X mental retardation protein (FMRP) in edited neurons (Liu et al., 2018). Another study finds that dCas9-DNMT3a target α-synuclein (*SNCA)* causes downregulation of SNCA mRNA and protein in patient derived dopaminergic neurons (Kantor et al., 2018). In cocaine exposure, epigenetic editing of histone methylation or acetylation at the FBJ osteosarcoma oncogene B (Fosb) locus using zinc finger protein is sufficient to dynamically regulate gene expression and control the behavioral effects of cocaine exposure (Heller et al., 2014). While showcasing the utility of epigenetic editing in brain, these studies have focused exclusively on regulating gene expression. The present study is the first to demonstrate the utility of epigenetic editing in regulating alternative splicing.

Finally, because global H3K36me3 enrichment regulated both cocaine reward behavior and alternative splicing of Srsf11-motif enriched genes, we hypothesized that Srsf11 is a key regulator of cocaine reward behavior. We therefore examined the effect of dCas9-Set2-mediated splicing of Srsf11 in cocaine CPP and found that this manipulation was sufficient to enhance drug reward behavior. Studies of splicing in other neurological conditions have highlighted the key role of isoform expression in mediating disease phenotypes. For example, one study in Alzheimer’s disease (AD) postmortem brains revealed that reduction in the ratio of transmembrane to soluble neurexin3 (NRXN3) promotes the neuronal AD phenotype (Hishimoto et al., 2019). Neurexin splicing variants also control postsynaptic response of N-methyl-D-aspartate (NMDA) and α-amino-3-hydroxy-5-methyl-4-isoxazolepropionic acid (AMPA) receptors. Specifically, alternative splicing of presynaptic neurexin-1 at splice site 4 (SS4) dramatically enhanced postsynaptic NMDA-receptor-mediated response, while alternative splicing of neurexin-3 at SS4 suppressed AMPA-receptor-mediated synaptic responses in hippocampus (Dai et al., 2019). These studies support the notion that alternative splicing can mediate different biological functions in varying neurological contexts. With this study, we provided additional evidence that alternative splicing events are associated with the pathology of addiction, such that Set2-mediated alternative splicing of Srsf11 is sufficient to augment cocaine preference.

### Conclusion

Set2 expression in NAc bidirectionally regulated cocaine preference. Epigenetic editing of H3K36me3 at Srsf11 was sufficient to drive intron retention of Srsf11 and enhance cocaine place preference and self-administration. We conclude that H3K36me3 functions directly in alternative splicing, and that this mechanism underlies cocaine reward behavior.

## METHODS

### Animals

Adult male, 8-week old C57BL/6J mice (Jackson) were used in this study. Mice were housed five per cage on a 12-h light-dark cycle at constant temperature (23°C) with access to food and water ad libitum. Animals were habituated to experimenter handling for at least 1 week before experimentation. All animal protocols were approved by the Institutional Animal Care and Use Committee of University of Pennsylvania.

### Intravenous cocaine self-administration (SA) and tissue collection

Mice were singly housed and habituated to the researcher by daily handling sessions for 7 days prior to the start of SA training. The acquisition of the instrumental task was facilitated in naïve mice by conducting daily training sessions in the operant chambers using a food pellet reward (20 mg). Mice were first introduced to the food pellets and food restricted (BioServ, Product #F0071), starting 3 days prior to the start of the operant training. Operant food training was conducted in 1-hour sessions for 10 days. The wheels were defined as <u>paired</u> if a 90° rotation of the wheel resulted in the presentation of a reward, with light and tone cues, and <u>unpaired</u> if spinning has no scheduled consequences. Following operant food SA training, mice were implanted with an indwelling catheter to the right external jugular vein under oxygen/isoflurane anesthesia and allowed 3 days to recover, during which time they were monitored daily for distress. Mice were then assigned to self-administer either saline or 0.56 mg/kg/infusion of cocaine on a fixed ratio (FR) schedule during daily 2-hour sessions for 21 consecutive days. Acquisition of cocaine SA was defined as three consecutive sessions during which ⩾ 10 infusions occurred, infusions did not vary by more than 20%, and at least 80% of spins were on the paired lever. For self-administration studies used for molecular biochemistry and sequencing, mice were run on a FR1 schedule for two-hour sessions over 21 consecutive days. For animals used in Set2-virus paired self-administration, an initial FR1 schedule was used. Once animals met learning criteria, they were moved to fixed-ratio 5 (FR5) and subsequently to progressive ratio (PR) schedule. 24 hours after the last cocaine session, mice were rapidly euthanized by cervical dislocation and decapitated, brains were removed and NAc, VTA, PFC was dissected using aluminum Harris micro-punch (Sigma-Aldrich). Tissue was frozen on dry ice and stored at −80 until downstream analysis.

### Cocaine conditioned place preference

Stereotactic surgery was performed as detailed below. After surgeries, mice were given 2 or 3 days to recover before being placed in the chamber and allowed to explore the conditioning apparatus. The conditioning is consisted of 2 distinct environment chambers (white/black stripe side, and gray side). Mice that showed pre-conditioning preference (more than 30% of total time spent in either of the 2 chambers) were excluded from the study. The initial preference was recorded (Pre-test). During training, mice received two parings per day: cocaine (15 mg/kg; i.p.) in the morning and were confined to the less preferred chamber in the pre-test. Saline (0.9%; 1 ml/kg; i.p.) in the afternoon and confined on the opposite side of the place preference chamber. On post-test day, mice were placed again in the either chamber with free access to both chambers, and the time spent in each side was quantified (Post-test). Data are expressed as percentage time spent on the cocaine-paired side in post-test minus the percentage time spent on the same side (CPP percentage time change).

### Factor Analysis

Factor analysis on cocaine SA was carried out based on published method (Walker et al., 2018). Briefly, A standard factor analysis was performed on cocaine SA behavioral variables using the scikit-learn package. This is to reduce the dimensions of behavioral variables in cocaine SA and generate a composite Addiction Index (AI) of 3 factors (Figure 3D, Supplement Figure 3F-L).

### Viral and Intra-NAc Transfection

Following anesthesia with a mixture of oxygen and isoflurane, 8-week-old C57BL/6 male mice were stereotactically infused with HSV virus encoding Set2 plasmid or catalytically dead Set2(R195G) control (Strahl et al., 2002). Nucleus accumbens (NAc) was targeted bilaterally using the following stereotaxic coordinates: + 1.6 (anterior/posterior), +1.5 (lateral), and −4.4 (dorsal/ventral) at an angle of 10° from the midline (relative to Bregma). The core and shell subregions of NAc are affected equally by these injections (the two subregions cannot reliably be targeted selectively in a mouse). A total of 1 μL of virus was delivered on each side over a 5-min period, followed by 5 min of rest. For Set2 with inhibitor Bay598 (SML1603) (10uM) treatment, inhibitor was dissolved in DMSO and mixed with virus. 1 μL of virus and inhibitor mixture was delivered to NAc on each side at a rate of 0.2 ul per minute, followed by 5 min of rest. dCas9-Set2 in-vivo transfection was conducted using the transfection reagent Jet-PEI (Polyplus transfection), prepared according to manufacturer’s instructions. 12.5 μl DNA plasmid (1.0 μg/uL) was diluted in 12.5 μl of sterile 10% glucose and added to diluted Jet-PEI, mixed by pipetting and incubated at room temperature (RT) for 15 min. A total of 1.5 μl of Jet-PEI/Plasmid solution was delivered NAc at a rate of 0.2 ul per minute, followed by 5 min of rest. In all molecular and behavioral experiments, proper NAc targeting of virus infusion was confirmed post hoc by preparing brain slices and visual confirmation of both needle track and GFP expression by microscopy. The mice were subjected to behavioral, immunohistochemical or DNA/RNA analysis after 2 days for virus transfection and 3 days for Jet-PEI transfection.

### Immunoblot analysis

Frozen NAc tissue were lysed in RIPA buffer (50 mM Tris-Cl, pH 8.0, 150 mM NaCl, 1% Nonidet P-40, 0.5% sodium deoxycholate and 0.1% SDS) plus a complete protease inhibitor cocktail (Roche). Lysates were centrifuged and supernatants were subjected to SDS-PAGE. Primary antibodies were as follows: rabbit polyclonal anti-H3K36me3 antibody (1:1,000, Abcam, #ab9050), mouse monoclonal anti-H3 antibody (1:2,000, Abcam, #ab24834). The blots were developed using an ECL kit (Pierce). Protein levels were quantified by ImageJ.

### Chromatin immunoprecipitation (ChIP) and ChIP-sequencing analysis

ChIP was performed on frozen bilateral NAc 2 mm punches pooled from 2-3 mice, and dissected as described above. Chromatin was prepared as described previously (Heller et al., 2014) and sheared using a diogenode bioruptor XL at high sonication intensity for 30 minutes (30 s on/30 s off). Fragment size was verified at 150–300 bp with an Agilent 2100 bioanalyzer. Sheared chromatin was incubated overnight with the H3K36me3 (Abcam ab9050) antibody previously bound to magnetic beads (Dynabeads M-280, Life Technologies. After reverse cross-linking and DNA purification (Qiagen Spin Column), ChIP-sequencing libraries were prepared using the NEBNext ChIP-Seq Library Prep Master Mix Set for Illumina (E6240L) using an adaptor oligo dilution of 1:20 for all samples. Prepared samples were pooled and sequenced on Illumina Hiseq 4000 platform to achieve ∼30 million reads per sample. Raw sequencing data was processed to generate fastq files of 50 bp single-end reads for further processing.

Sequences were aligned using Bowtie v2.1.0, allowing for ⩽2 mismatches to the reference in 50 bp. High quality and uniquely aligned reads were confirmed for H3K36me3 ChIP-sequencing. Uniquely mapped reads were used for subsequent analysis. Peak calling was performed using SICER (Xu et al., 2014), whereas DiffBind (Ross-Innes et al., 2012) was used to identify the differential histone modification sites between treatment and control groups.

Integrated profiles were created by averaging the observed signal for each bin for the selected set of relevant genes using deepTools computeMatrix; values displayed indicate fragments per 50-bp bin per million mapped reads (Figures 1H, 2H, 5B-D, Supplement figure 2C). Integrated profile of H3k36me3 enrichment at splice junctions (Figure 1G, 4C, Supplement figure 4C) was created similarly using deepTools computeMatrix. Splice junctions were defined using 200bp up- and down-stream of spliced exon start/end site. Permuted splicing junctions were generated by randomly permuting the genomic locations of the splice junctions using Bedtools shuffle. H3K36me3 Enrichment of islands over genome was assessed using HOMER software with a false discovery rate (FDR) of 5% as a cutoff and a significance score (fold-change score over input) of 1.5. By default, HOMER used DESeq2’s rlog transform to create normalized log2 read counts, thereafter generate log2 transformed enrichment relative to a specific genomic category (Heinz et al., 2010). Gene Ontology was analyzed using the Database for Annotation, Visualization and Integrated Discovery (DAVID) Bioinformatics Resource specifying for biological processes. IGV is used to visualize the track coverage.

### RNA extraction, PCR and quantitative RT-PCR

Total RNA was extracted from tissue and cells using the RNeasy Mini Kit (Qiagen) as recommended by the manufacturer. To assess inclusion of alternative exons, forward and reverse primers were designed to anneal to the constitutively included exons upstream and downstream of the alternative exon, respectively. PCR assays were performed based on previous published method (Gonatopoulos-Pournatzis et al., 2018) OneStep RT-PCR kit (QIAGEN) was used according to the manufacturer’s recommendations. Reaction products were separated on 1% agarose gels. Percent Spliced In (PSI) values were calculated using ImageJ software. First the exon-included and exon-excluded band intensities were corrected by subtracting background. Then, intensity of the exon-included band was divided by the sum of the exon-included and exon-excluded bands. The result was multiplied by 100% to obtain the PSI value. qPCR and data analysis was performed as previously described (Heller et al., 2016), by comparing Ct values of the experimental group to control using the ΔΔCt method.

### RNA-sequencing and analysis

RNA quality control assays and library preparation were performed by the University of Pennsylvania Next Generation Sequencing core. Total RNA was quantified using fluorescent chemistry contained in the Qubit RNA HS Assay Kit (Cat# Q10211, Thermo Fisher Scientific). 1 ul was used to assess RNA Integrity Number (RIN) using the Agilent Bioanalyzer RNA nano kit (Cat # 5067-1513, Agilent). The RIN range was between 8 and 9.8. RNA-seq was carried out on an Illumina HiSeq 4000. Sequence alignment was performed using STAR (Dobin et al., 2013). The FASTQ sequence was aligned to the mouse genome mm9 or mm10 and unique read alignments were used to quantify expression and aggregated on a per-gene basis using the Ensembl (GRCm38.67) annotation. We used DESeq2 (Love et al., 2014) to assess differential expression between sample groups, and both rMATS (Shen et al., 2014) and MAJIQ (Green et al., 2017; Norton et al., 2018) to assess alternative splicing. Cocaine SA splicing data (Figure 4A-B) was defined as MAJIQ identified alternative spliced genes common to cocaine SA plus 1-day and 28-day abstinence. All RNA-seq data was deposited in NCBI as GEOxx.

### Extraction of histones from NAc tissue

Tissue was lysed in nuclear isolation buffer (15 mM Tris pH 7.5, 60 mM KCl, 15 mM NaCl, 5 mM MgCl2, 1 mM CaCl2, 250 mM sucrose, 10 mM sodium butyrate, 1 mM DTT, 500 µM AEBSF, 5 nM microcystin) containing 0.3% NP-40 alternative on ice for 5 min. Nuclei were pelleted and resuspended in 0.4 N H2SO4 followed by 1.5 hr rotation at 4°C. After centrifugation, supernatants were collected, proteins were precipitated in 33% TCA overnight on ice, washed with acetone, and resuspended in deionized water. Acid-extracted histones (5-10 μg) were resuspended in 100 mM ammonium bicarbonate (pH 8), derivatized using propionic anhydride and digested with trypsin as previously described (Sidoli et al., 2016). After a second round of propionylation the resulting histone peptides were desalted using C18 Stage Tips, dried using a centrifugal evaporator, and reconstituted using 0.1% formic acid in preparation for LC-MS analysis.

### Liquid Chromatography – Mass Spectrometry

Nanoflow liquid chromatography was performed using a Thermo ScientificTM Easy nLCTM 1000 equipped with a 75 µm x 20 cm column packed in-house using Reprosil-Pur C18-AQ (3 µm; Dr. Maisch GmbH, Germany). Buffer A was 0.1% formic acid and Buffer B was 0.1% formic acid in 80% acetonitrile. Peptides were resolved using a two-step linear gradient from 5% to 33% B over 45 min, then from 33% B to 90% B over 10 min at a flow rate of 300 nL/min. The HPLC was coupled online to an Orbitrap Elite mass spectrometer operating in the positive mode using a Nanospray Flex™ Ion Source (Thermo Scientific) at 2.3 kV.

For MS, a full scan (m/z 300-1100) was acquired in the Orbitrap mass analyzer with a resolution of 120,000 (at 200 m/z) followed by 8 MS2 scans using precursor isolation windows of 50 m/z each (e.g. 300-350, 350-400…650-700). A second full scan was then performed followed by the remaining 8 MS2 scans (800-850, 900-950…). MS/MS spectra were acquired in the ion trap operating in normal mode. Fragmentation was performed using collision-induced dissociation (CID) in the ion trap mass analyzer with a normalized collision energy of 35. AGC target and maximum injection time were 5e5 and 50 ms for the full MS scan, and 3e4 and 50 ms for the MS2 scans, respectively. Raw files were analyzed using EpiProfile 2.0 (Yuan et al., 2018). The area under the curve (AUC) for each modified state of a peptide was normalized against the total of all peptide AUCs to give the relative abundance of the histone modification.

### Plasmid construction and sgRNA design

The Set2 and Set2 mutant R195G plasmid used for this study were obtained from Dr. Brian Strahl (UNC). Plasmids were then packaged into HSV-GFP virus. dCas9-Set2 construct was optimized, synthesized and cloned into pENTR-dTOPO by GENEWIZ. Then the entry plasmid pENTR-dTOPO-dCas9-Set2 was packaged into destination vector hSyn-GW-IRES2-mCherry using Gateway LR Clonase II enzyme mix (ThermoFisher 11791020). The hsyn-GW-dCas9-SET2-IRES2-mCherry vector, encoding mammalian codon-optimized Streptococcus pyogenes dCas9 fused to a human SET2 domain were used for neuronal cell line and neuronal tissue transfections.

The single guide RNA was designed and screened using the web tool http://crispr.mit.edu/. The PAM sequence of these guide RNAs was used to determine the minimal off-target effect. The three sgRNAs with the highest off-target scores were designed based on H3K36me3 enriched region and strand specificity. Then the 19bp target sequence was synthesized into gBlock that contains U6 promoter, guide RNA scaffold and termination signal. Then gBLOCK was cloned into an empty backbone vector pENTR/d-TOPO (ThermoFisher K240020) following manufacture instructions. Additionally, the entry plasmid was packaged into destination vector P1005-GFP using Gateway LR Clonase II enzyme mix (ThermoFisher 11791020).

### Cell culture and transfection

Neuro2a cells (CCL-131), ATCC were cultured in EMEM medium with 10%FBS in 6-well plates. The cells were transfected with 400 ng of plasmid DNA using Effectene reagent (Qiagen), and RNA was isolated using the RNeasy Mini Kit (Qiagen) according to the manufacturer instructions. The cells were maintained at 37 °C and 5% CO2, and harvested in 24 hours. dCas9-Set2 with sgRNA were transfected in six-well plates with 1 mg of each respective dCas9 expression vector, 0.33 mg of individual gRNA expression vectors using Effectene reagent (Qiagen) as per the manufacturer’s instruction. The cells were maintained at 37 °C and 5% CO^2^, and harvested in 48 hours after transfection.

### Splicing motif analysis

De novo motifs (recurring, fixed-length patterns) in the top 1000 (rank by “inter” treatment difference) alternative splicing junctions (+-200bps of splice site) were identified using HOMER software with scrambled sequence as control. Similar motif analysis was done using MEME. Searched motifs were then submitted as input to the MEME-Tomtom program to rank the motifs in the database and produce an alignment for each significant match for splice factors using Ray2013 RBP databases (Ray et al., 2013).

### CUT&RUN sequencing

CUT&RUN was performed based on published method (Skene and Henikoff, 2017; Skene et al., 2018). Transfected N2a cells were first lysed to isolate the nuclei. The nuclei were then centrifuged, washed and incubated with lectin-coated magnetic beads. The lectin-nuclei complexes were then resuspended with anti-H3K36me3 antibody (Abcam, #ab9050) and incubated overnight at 4°C. The nuclei were washed next day to remove unbound antibodies and incubated with Protein-A-MNase for 1 hour, then washed again to remove unbound protein-A-MNase. CaCl2 was added to initiate the calcium dependent nuclease activity of MNase to cleave the DNA around the DNA-binding protein. The protein-A-MNase reaction was quenched by adding chelating agents (EDTA and EGTA). The cleaved DNA fragments were then liberated into the supernatant and incubated with proteinase K for 15 minutes at 70°C. DNA fragments were purified using Qiagen mini-elute and used to construct a sequencing library using NEBNext ChIP-Seq Library Prep Master Mix Set for Illumina® (E6240L) using an adaptor oligo dilution of 1:20 for all samples. Samples were pooled and sequenced on Illumina Hiseq 4000 platform to achieve ∼20 million reads per sample. Raw sequencing data was processed to generate fastq files of 50 bp pair-end reads for further processing. The data was then collected and analyzed using software that aligns sample sequences to mm9 to identify the H3K36me3 targeted DNA fragments.

## ACKNOWLEDGEMENTS

The authors wish to thank Dr. Brian D Strahl (University of North Carolina) for generously donating the Set2 and Set2(R195G) plasmids used in this study. The authors thank Dr. Yoseph Barash and Matthew Gazzara for assistance in using MAJIQ. The authors thank Dr. Erica Korb (University of Pennsylvania) for critical reading of the manuscript. Financial support is kindly acknowledged from Charles E Kaufman Foundation Young Investigator Award (E.A.H), Whitehall Foundation Grant (E.A.H), NIH-NIDA Avenir Director’s Pioneer Award (E.A.H, DP1 DA044250) and Research Supplements to Promote Diversity in Health-Related Research (M.D.C, DP1 DA044250-01), and T32 Predoctoral Training Grant in Pharmacology (M.D.C, T32GM008076). The cocaine used in this study was kindly provided by the NIDA drug supply program.

## SUPPLEMENT FIGURE LEGENDS

**Supplement figure 1.**
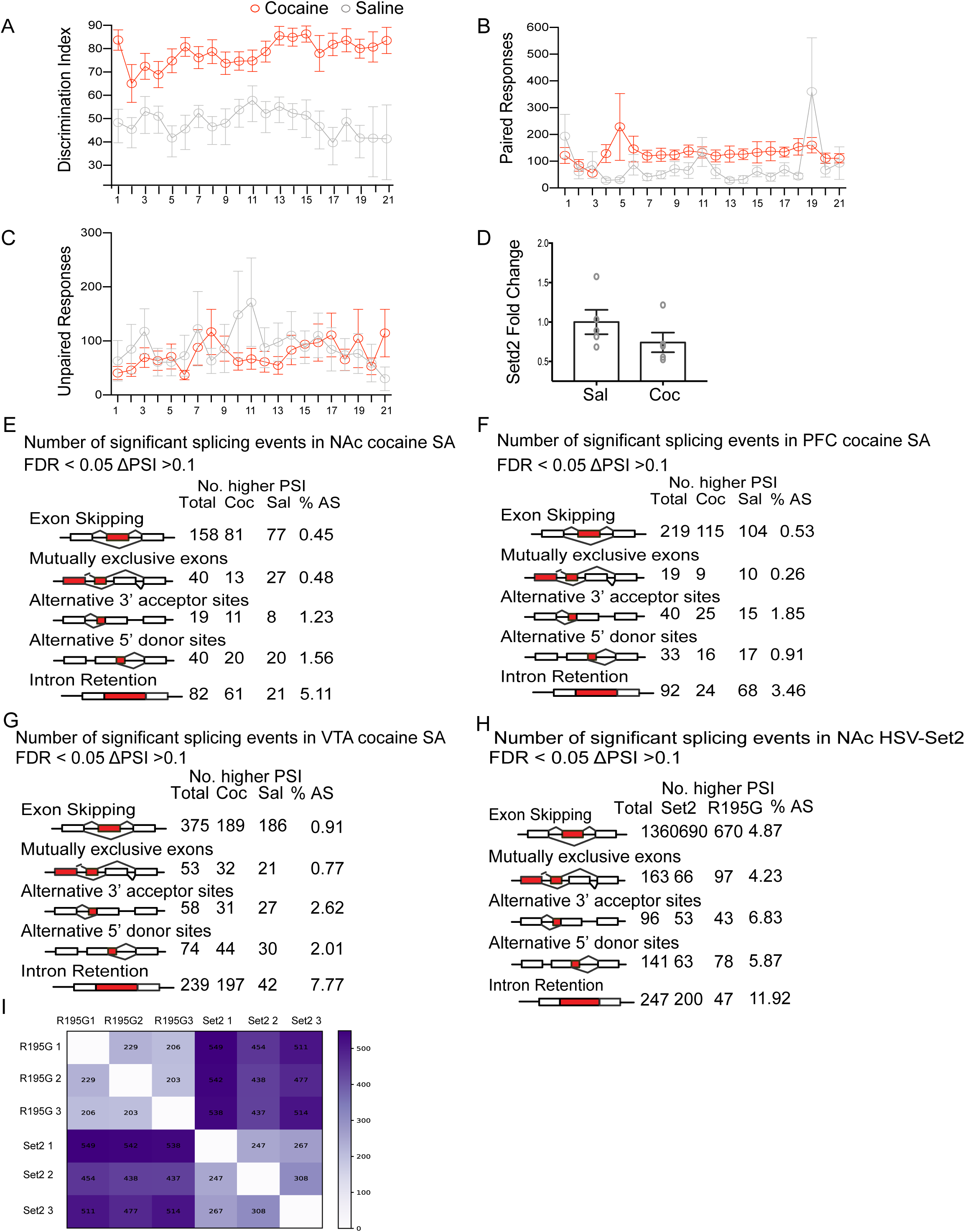
Cocaine SA behavior and alternative splicing profiles in multiple brain regions. (A) Mice show significantly greater discrimination between the paired and unpaired wheel sessions (two-way ANOVA repeated measure). Interaction effect F(20,440) = 1.002, P=0.4584; drug effect F(1,22)=36, P<0.0001; session effect F(24,440) =1.411, P=0.1116). (B) Mice in cocaine group spun the paired wheel significantly more than saline group across sessions (two-way ANOVA repeated measure). Interaction effect F(20,440) = 1.487, P =0.0810; drug effect F(1,22)=9.406, P=0.0056; session effect F(24,440) =0.9203, P=0.5612). (C) Cocaine-SA mice spun the unpaired wheel significantly less than saline-SA mice across sessions (two-way ANOVA repeated measure; Interaction effect F(20,440) = 1.407, P =0.1136; drug effect F(1,22)=0.07903, P=0.7812; session effect F(24,440) =0.9474, P=0.5266). (D) Biological validation of Setd2 expression levels in NAc following cocaine SA, compared to saline controls. (E) Positive linear correlation between H3K36me3 enrichment (western blot, NAc) and cocaine CPP. HSV treatment groups include intracranial injection of Set2, Set2(R195G), Set2+Inhibitor and Set2(R195G)+Inhibitor (R^2^ = 0.42, P<0.05). (F-H) Alternative splicing summary comparing SA cocaine and saline in (E) NAc, (F) PFC and (G) VTA identified by rMATS. Only alternative splicing events that are FDR < 0.05 and ΔPSI > 0.1 were shown in the table. (I) Alternative splicing summary comparing HSV-Set2 and HSV-Set2(R195G) in NAc identified by rMATS. Only alternative splicing events that are FDR < 0.05 and ΔPSI > 0.1 were shown in the table. (J) Alternative splicing events summary comparing HSV-Set2 and HSV-R195G in NAc identified by MAJIQ V1.1 using pairwise comparison of each replicate. Most of alternative splicing events occurred between treatment in the pairwise comparison as indicated by the dark color. Alternative splicing events are also detected within treatment due to sample to sample variance.

**Supplement figure 2.**
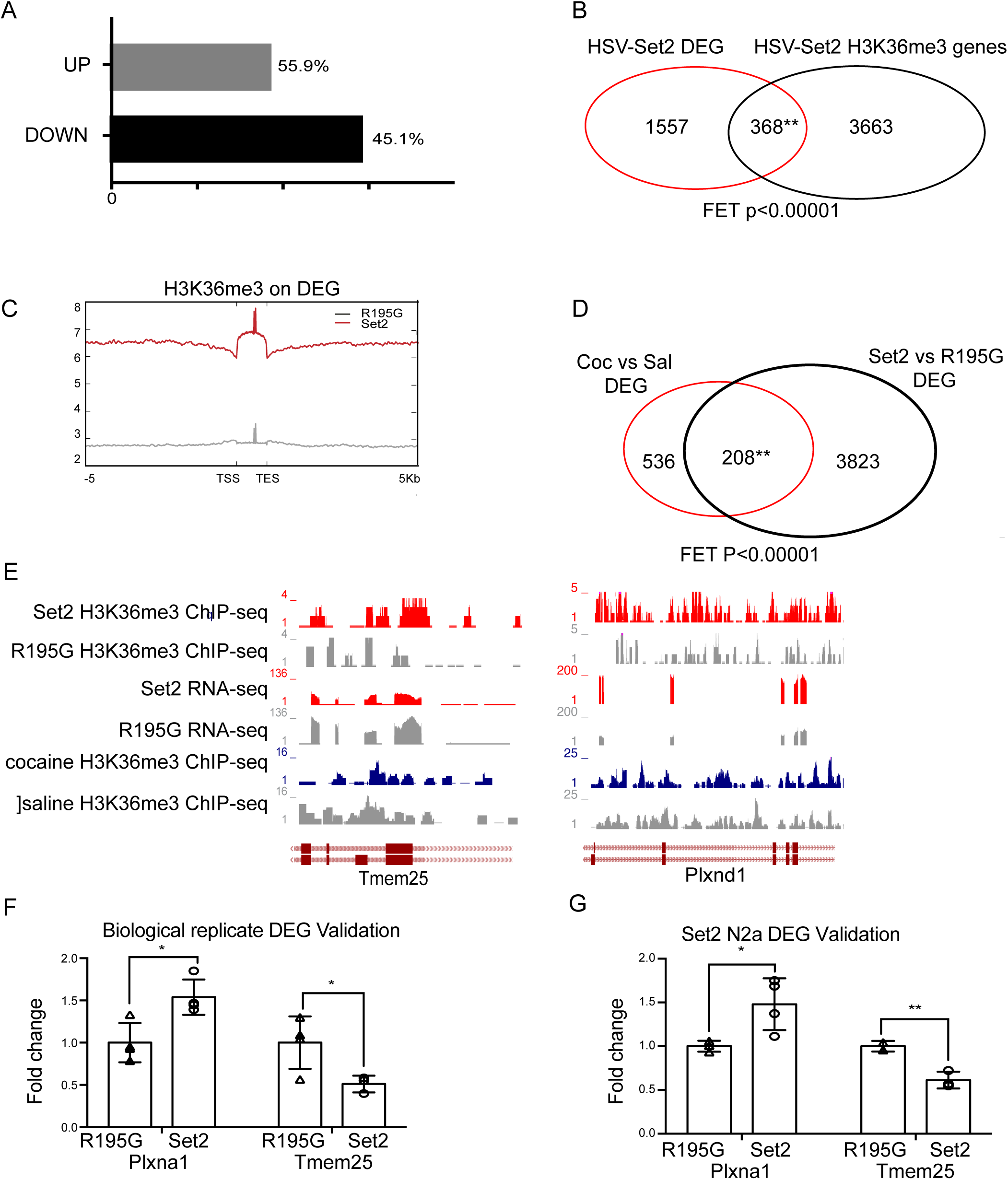
Set2 overexpression regulates gene expression in NAc. (A) HSV-Set2 up- and down-regulates gene expression. Among the regulated genes, 55.9% were upregulated comparing to HSV-Set2(R195G) and 45.1% were down-regulated. (B) Venn diagram showing overlap between DEG identified by RNA-seq and HSV-Set2 mediated H3K36me3 enriched genes (FET < 0.00001). (C) Profile plot of H3K36me3 enrichment on HSV-Set2 mediated DEGs compared to Set2(R195G). (D) Venn diagram showing overlap between DEGs mediated by cocaine compared to saline and Set2 compared to Set2(R195G) (FET< 0.00001). (E) Track coverage of H3K36me3 ChIP- and RNA-seq from HSV-Set2 and -Set2(R195G), as well as cocaine and saline for representative up-regulated gene *Tmem25* and down-regulated gene *Plxnd1*. (F) qPCR validation on Set2-mediated DEGs from separate biological replicates. (G) qPCR validation on Set2-mediated DEGs from transfected Neuro2a cells.

**Supplement figure 3.**
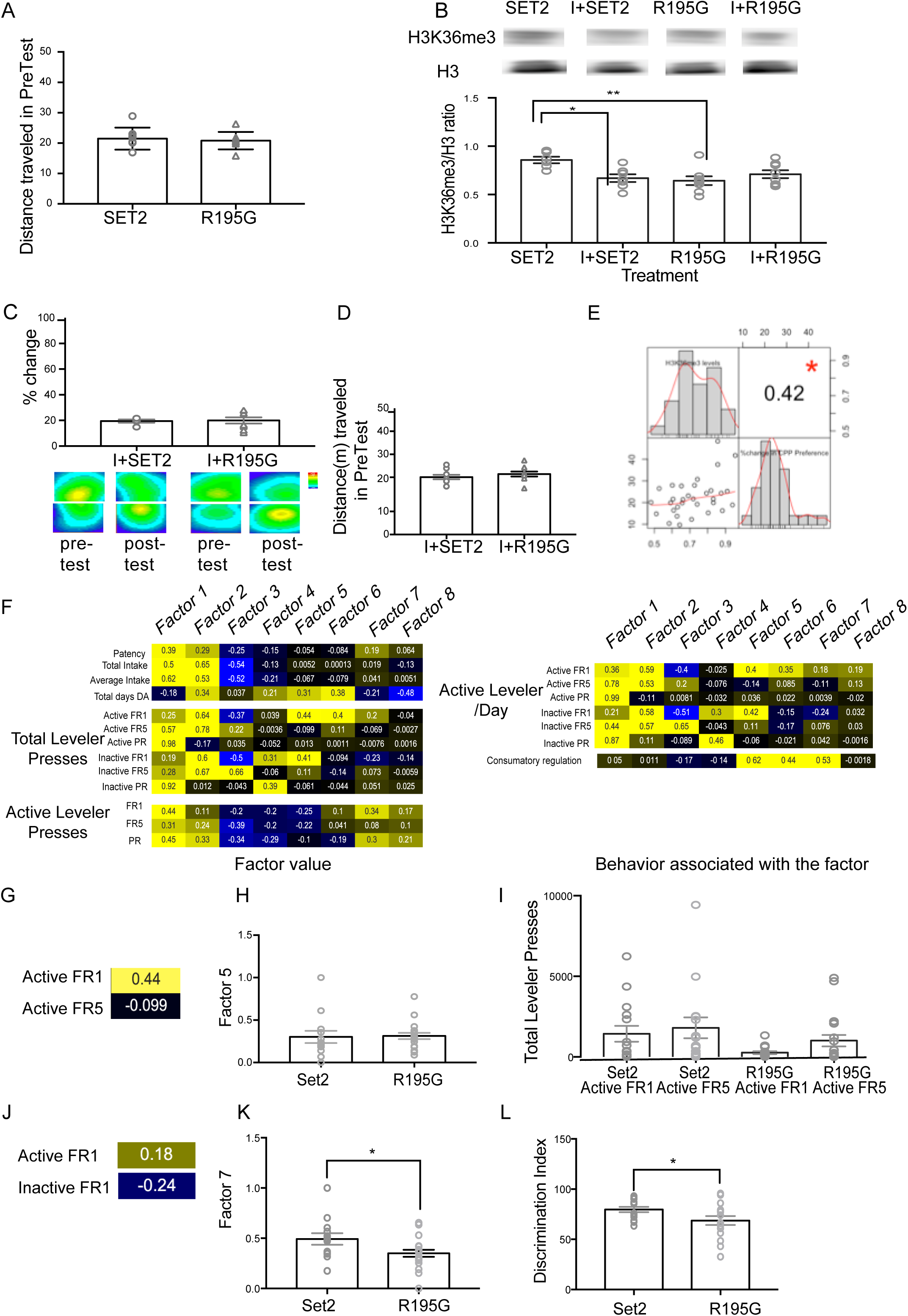
Set2 mediated H3K36me3 enrichment increases cocaine preference. (A) Distance traveled in pre-test by HSV-Set2 and HSV-Set2(R195G) injected animals (non-significant, Student’s unpaired t-test). (B) Representative western blot for H3K36me3 and total H3 in HSV-Set2, HSV-Set2 with inhibitor Bay598, HSV-R195G, and HSV-R195G with methyltransferase inhibitor Bay598 injected NAc (one-way ANOVA with Tukey’s multiple comparison, * P<0.05,** P<0.01). BAY598 is a methyltransferase inhibitor that reduces the effect of SET2 on CPP. This confirmed that the Set2-mediated increase in cocaine CPP required SET2 catalytic activity. (C) Percentage change in time spent in cocaine treated side of HSV-Set2 and HSV-Set2 (R195G) with BAY598 injected animals. (D) Distance traveled in Pre-Tests of HSV-Set2 and HSV-Set2(R195G) with BAY598 injected animals. (E) Factor analysis was used to reduce multidimensional behavioral data to factors. The association of each factor with each behavioral endpoint included in the analysis is displayed. Factors were positively (yellow), negatively (blue), or not associated (black) with each endpoint. These particular associations allowed for the interpretation of the how each factor related to Set2 affected cocaine SA behaviors. (F-K) Data for individual animals for each behavior and each factor are presented. (F, I) Factor loading of factors 5 and 7 with self-administration behaviors (yellow = positive; blue = negative) are presented. (G, J) Individual data presented for the behaviors associated with each factor. (G) Factor 5 is associated with consummatory regulation. It positively associated with paired lever under a fixed-ratio 1 and negatively associated with unpaired lever under a fixed-ratio 5, and (J) Factor 7 is positively associated with paired lever and negatively associated with unpaired lever under a fixed-ratio 1. (H, K) Individual transformed data for (H) factor 5 associated total levels presses in Set2 or Set2(R195G) in cocaine SA experiment (Student T-test, * P<0.05), and (K) factor 7 associated discrimination index in Set2 or Set2(R195G) in cocaine SA experiment (Student T-test, * P<0.05).

**Supplement figure 4.**
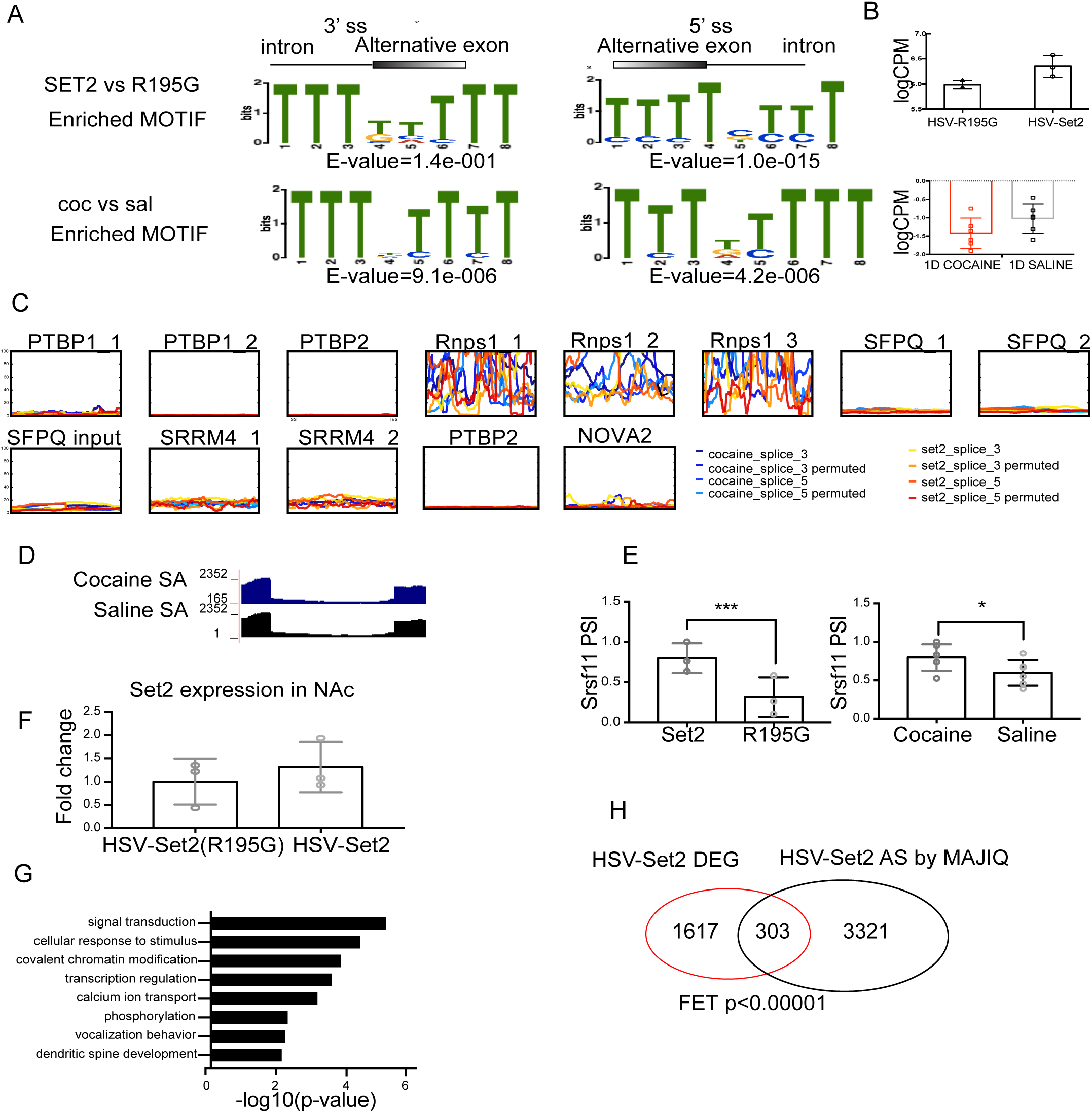
Srsf11 is identified as a key splice factor that mediates alternative splicing in cocaine and Set2 treatments. (A) Representative splicing motifs in cocaine SA and Set2 at both 3’ and 5’ splice junctions that were identified by Meme. (B) Expression of Srsf11 in cocaine SA and Set2 detected in RNA-seq. No significant differential expression was identified in either comparison. (C) Published CLIP-Seq signal plotted on cocaine SA and Set2 splicing junctions and their corresponding permuted sequences at both 3’ and 5’. None of the signal showed a distinct pattern such as Srsf11. (D) Track coverage of RNA-seq of cocaine and saline samples in cocaine SA. (E) Srsf11 alternative splicing event was detected by rMATs in cocaine SA and Set2 (* FDR<0.05, **FDR<0.01). (F) Validation on Srsf11 expression of Set2 and Set2(R195G) controls using biological replicates (No significance, Student T-test). (G) Function analysis on overlapped alternative splicing genes among cocaine SA and Set2 group. (H) Venn diagram showing significant overlap between DEGs and alternative splicing genes mediated by Set2 overexpression (FET P <0.00001).

**Supplement figure 5.**
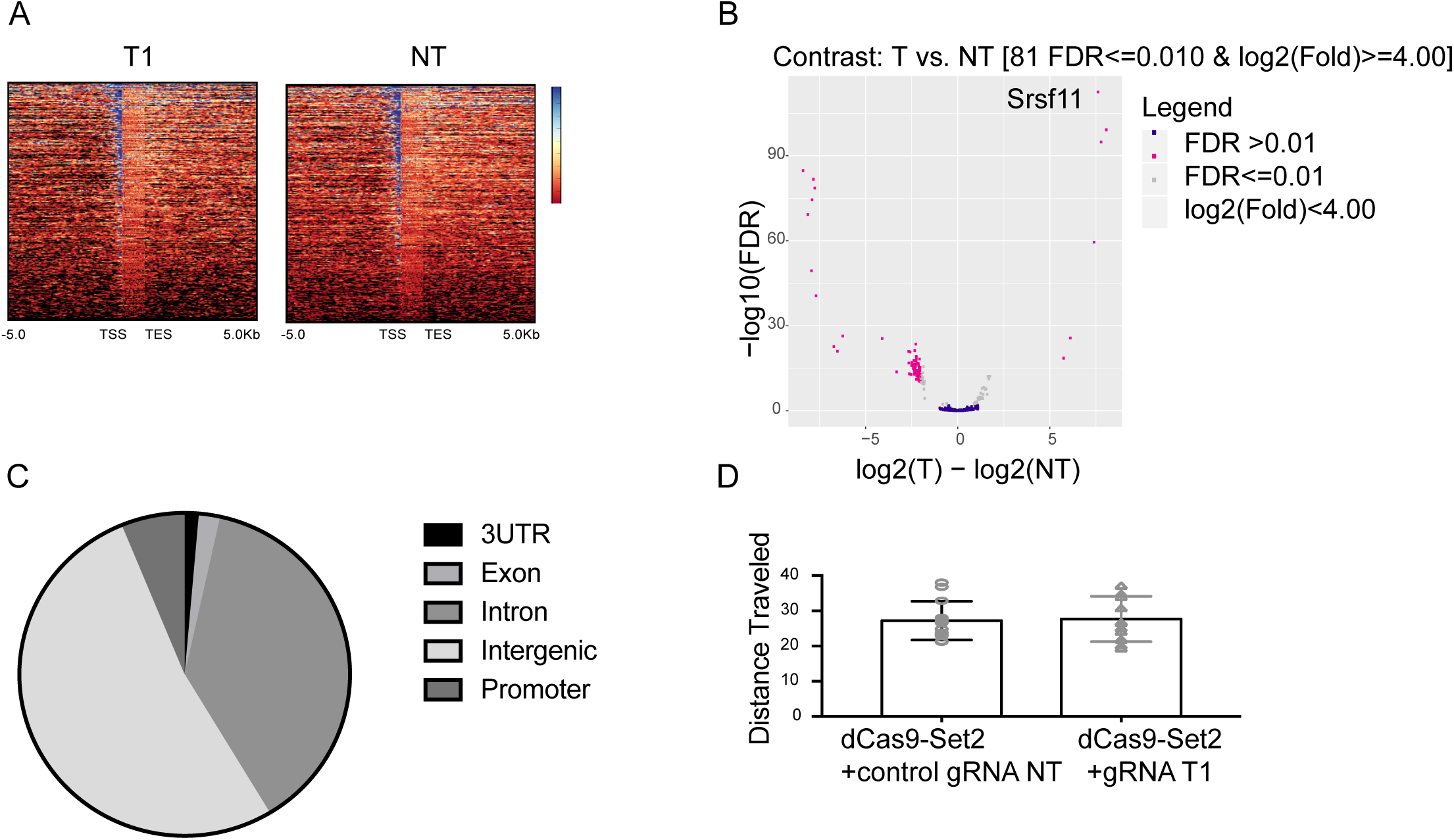
dCas9-Set2 shows negligible off-target effect in mediating H3K36me3 enrichment. (A) Heatmap of genome-wide H3K36me3 enrichment of dCas9-Set2 plus sgRNA T1 and NT transfected N2a cells identified by CUT&RUN-seq. No difference in enrichment distribution was observed between the two treatments. (B) Volcano plot from DiffBind analysis between dCas9-Set2 with sgRNA T1 and NT. Srsf11 was the highest differentially enriched gene (Fold change = 152.2, FDR = 5e-75). Pink dots indicate other significantly differential H3K36me3 enriched peaks (log2(Fold-change)>=4). (C) Genomic locations of 122 identified H3K36me3 differential enrichment in dCas9-Set2 plus sgRNA T1 compared to dCas9-Set2 plus sgRNA NT. The majority of off-target enrichment are at intergenic regions. (D) Distance traveled in pre-test by dCas9-Set2 plus sgRNA T1 compared to dCas9-Set2 plus sgRNA NT animals (non-significant, Student’s unpaired t-test).

